# Mechanotransduction and dynamic outflow regulation in trabecular meshwork requires Piezo1 channels

**DOI:** 10.1101/2020.06.30.180653

**Authors:** Oleg Yarishkin, Tam T. T. Phuong, Jackson M. Baumann, Michael L. De Ieso, Felix Vazquez-Chona, Christopher N. Rudzitis, Chad Sundberg, Monika Lakk, W. Daniel Stamer, David Križaj

**Affiliations:** Department of Ophthalmology and Visual Sciences, University of Utah, Salt Lake City, UT 84132, USA; Department of Bioengineering, University of Utah, Salt Lake City, UT 84132, USA; Duke Eye Center, Duke University School of Medicine, Durham, NC 27710, USA; Department of Neurobiology and Anatomy, University of Utah School of Medicine, Salt Lake City, UT 84132, USA

**Keywords:** Trabecular meshwork, mechanosensation, shear stress, ion channel, Piezo1

## Abstract

Mechanosensitivity of the trabecular meshwork (TM) is a key determinant of intraocular pressure (IOP) yet our understanding of the molecular mechanisms that subserve it remains in its infancy. Here, we show that mechanosensitive Piezo1 channels modulate the TM pressure response via calcium signaling and dynamics of the conventional outflow pathway. Pressure steps evoked fast, inactivating cation currents and calcium signals that were inhibited by Ruthenium Red, GsMTx4 and Piezo1 shRNA. Piezo1 expression was confirmed by transcript and protein analysis, and by visualizing Yoda1-mediated currents and [Ca^2+^]_i_ elevations in primary human TM cells. Piezo1 activation was obligatory for transduction of physiological shear stress and was coupled to reorganization of F-actin cytoskeleton and focal adhesions. The importance of Piezo1 channels as pressure sensors was shown by the GsMTx4 -dependence of the pressure-evoked current and conventional outflow function. We also demonstrate that Piezo1 collaborates with the stretch-activated TRPV4 channel, which mediated slow, delayed currents to pressure steps. Collectively, these results suggest that TM mechanosensitivity utilizes kinetically, regulatory and functionally distinct pressure transducers to inform the cells about force-sensing contexts. Piezo1-dependent control of shear flow sensing, calcium homeostasis, cytoskeletal dynamics and pressure-dependent outflow suggests a novel potential therapeutic target for treating glaucoma.

**Significance Statement:** Trabecular meshwork (TM) is a highly mechanosensitive tissue in the eye that regulates intraocular pressure through the control of aqueous humor drainage. Its dysfunction underlies the progression of glaucoma but neither the mechanisms through which TM cells sense pressure nor their role in aqueous humor outflow are understood at the molecular level. We identified the Piezo1 channel as a key TM transducer of tensile stretch, shear flow and pressure. Its activation resulted in intracellular signals that altered organization of the cytoskeleton and cell-extracellular matrix contacts, and modulated the trabecular component of aqueous outflow whereas another channel, TRPV4, mediated a delayed mechanoresponse. These findings provide a new mechanistic framework for trabecular mechanotransduction and its role in the regulation of fast fluctuations in ocular pressure, as well as chronic remodeling of TM architecture that epitomizes glaucoma.

## Introduction

Maintenance of hydrostatic pressure gradients in the body requires active mechanosensing by nonexcitable cells. Paradigmatic examples from organs that experience high dynamic mechanical loads include the myogenic response that protects the brain, systemic vasculature and lung from hypertension-induced injury (1, 2) control of water/solute flow in the kidney and bladder (3), and the ocular outflow system, which responds to fluctuations in intraocular pressure (IOP) with compensatory adjustment of aqueous humor drainage via the conventional (trabecular) outflow pathway (4-7). Outflow homeostasis depends in part on mechanotransduction and contractility of trabecular meshwork (TM) cells, which dynamically regulate the tissue resistance to fluid flow (8). In ocular hypertensive glaucoma, the irreversible remodeling of TM architecture (9, 10) increases the risk for developing optic neuropathy (11, 12) whereas *in vitro* studies show upregulation of glaucoma-associated genes upon exposure of TM cells to cyclic strains (5, 13). A key missing link, however, remains the lack of understanding of the mechanisms that convert mechanical stimuli into intracellular signals that regulate trabecular remodeling and thus outflow.

Pressure homeostasis in the vasculature, lung, bladder and eye is dynamically regulated by stretch-activated ion channels (SACs) (2, 14, 15) that belong to Piezo, TRP (transient receptor potential) and two-pore potassium channel families (16). TRPV4, a polymodal nonselective cation channel, mediates the TM Ca^2+^ influx in response to stretch and contributes to cell remodeling in the presence of chronic pressure/strain (15, 17) but the mechanotransduction mechanism that mediates the homeostatic adjustment to pressure fluctuations has remained unknown. Potential candidates are Piezo channels, which have been recently implicated in dynamic regulation of pressure-induced lung vascular hyperpermeability and shear-stress induced endothelial signaling (18, 19). Piezo 1 and 2 are large, conserved trimeric channels that mediate rapidly inactivating, small conductance (∼25 - 35 pS), low-threshold (∼0.6 kBT/nm^2^) currents in response to membrane tension (∼1 - 3 mN/m), indentation, shear stress (∼10 dyn/cm^2^) and/or stretch (20-23). The Genotype-Tissue Expression (GTEx) project (24) shows Piezo1 expression in nonexcitable cells from several pressure regulating tissues, and the channel was shown to play a central role in the regulation of cell volume, heart rate and flow-mediated vasoconstriction by smooth muscle cells (SMCs) and endothelial cells (25-29) but its roles in ocular tissues are virtually unknown.

In this study, we show Piezo1 is strongly expressed in human and mouse TM, identified it as a principal transducer of membrane tension and shear flow and linked its activation to dynamic control of the TM cytoskeleton, cell-ECM contacts and aqueous outflow. Whereas TRPV4 channels mediated a delayed component of the stretch-activated current. These findings suggest that TM mechanotransduction involves contributions from kinetically and pharmacologically distinguishable SACs. Our data also suggest a homeostatic function for Piezo1-dependent trabecular outflow in response to dynamic changes in pressure-induced stretch, with potential implications for the etiology and treatment of glaucoma.

## Matherials and Methods

See SI Experimental Procedures for more details

### Animals

C57BL/6J and *Piezo1*^*P1-tdT*^ (*Piezo1*^*tm1*.*1Apat*^) mice were from JAX Labs. The initial transgenic 129/SvJ strain (30) was backcrossed to C57BL/6J for more than 8 generations. The animals were maintained in a pathogen-free facility with a 12-hour light/dark cycle and *ad libitum* access to food and water.

### Cell culture

TM cells were isolated from juxtacanalicular and corneoscleral regions of the human donors’ eyes, as described (15, 17), in accordance with consensus characterization recommendations (31).

### Reagents

Reagents used were largely purchased from Sigma-Aldrich. *Grammostola spatulata* mechanotoxin 4 (GsMTx4) were from Sigma-Aldrich and Alomone Labs.

### Western Blots

Total protein samples were extracted from two primary trabecular meshwork lines obtained from different patients in completed RIPA buffer supplemented with an enzyme inhibition cocktail (Biotechnology, Inc., Santa Cruz, CA, USA). Protein concentration was estimated with the Bradford assay.

### Immunohistochemistry

Anterior chambers were fixed in 4% para-formaldehyde for one hour, cryoprotected in 15 and 30% sucrose gradients, embedded in Tissue-Tek® O.C.T. (Sakura, 4583), and cryosectioned at 12 µm, as described (32) (33). The sections were probed with antibodies against Piezo1 (Proteintech, 15939), α-SMA (Sigma, A2457), and Collagen IV (EMD Millipore, AB769). Secondary antibodies included anti-rabbit IgG DyLight 488 (Invitrogen, 35552), anti-mouse IgG DyLight 594 (Invitrogen, 35511), and anti-goat IgG Alexa 647 (Invitrogen, A21469).

### Calcium imaging

Patch clamp data was acquired with a Multiclamp 700B amplifier, pClamp 10.6 software and Digidata 1440A interface (all from Molecular Devices). Data was sampled at 5 kH, digitized at 2 kH and analyzed with Clampfit 10.7 (Molecular Devices) and Origin 8 Pro (Origin lab). Steps of positive (in whole-cell recording) and negative (in single-channel recording) pressure were delivered via High-Speed Pressure Clamp (ALA Scientific) as described (**Figure 1A)** (17, 34).

**Figure 1.**
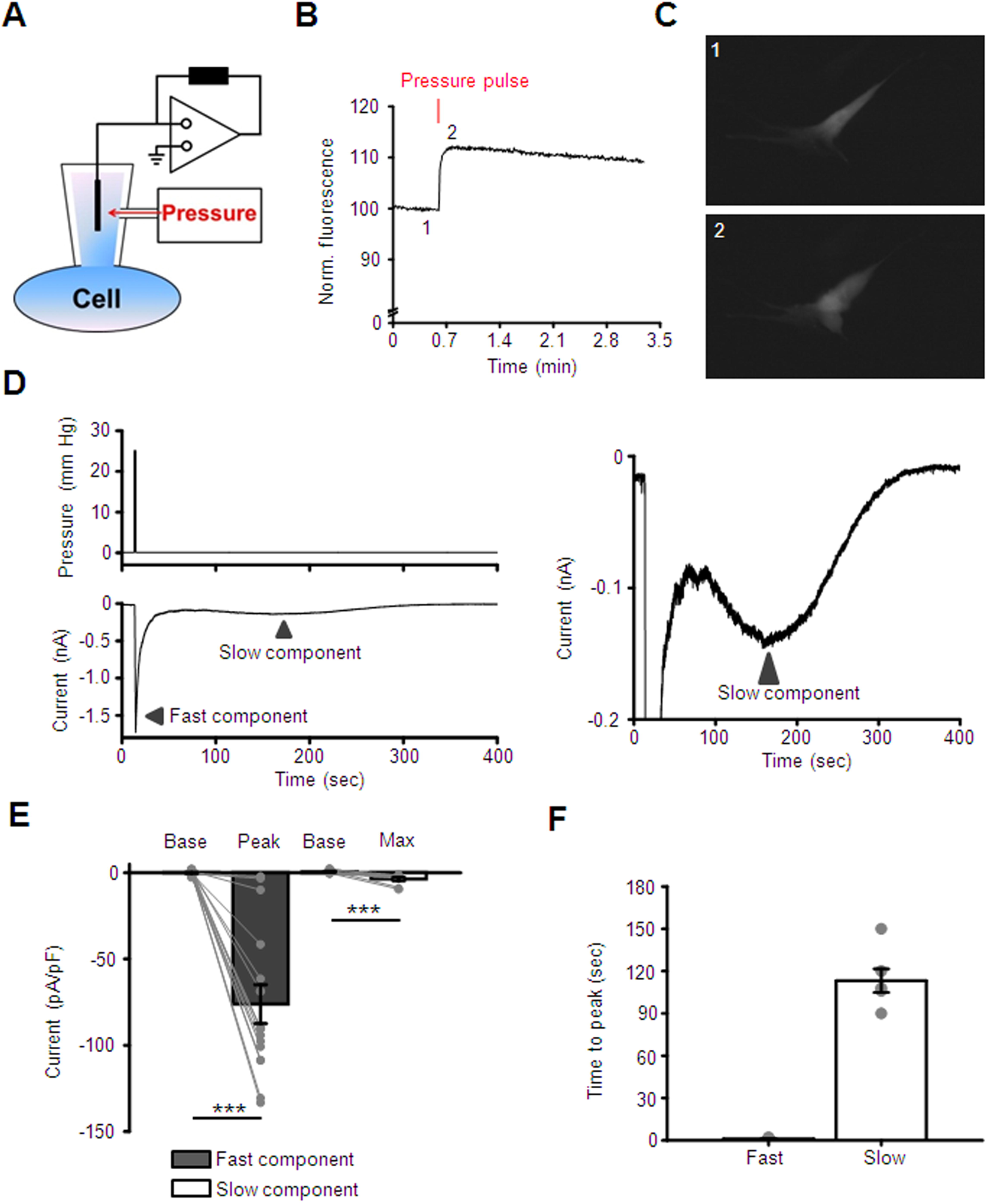
Piezo1 and TRPV4 mediate pressure-induced current of distinct kinetics in TM cells. **(A)** Schematic diagram of the setup. **(B)** A representative trace of calcein fluorescence illustrating effect of pressure pulse. Positive changes in fluorescence indicate at increase in cell volume. **(C)** A representative fluorescent image demonstrating change in cell volume after application of a pressure pulse. Images were taken at corresponding time points shown in (B). **(D)** Representative trace of the fast and the slow components *(lower trace)* of transmembrane current evoked by pressure pulse *(upper trace)*. The right panel is the Y-axis expansion of the trace shown in the left panel. The holding potential was -40 mV. **(E)** Quantification of the magnitude of the fast and the slow responses. Shown are the means ± SEM of baseline current (base) measured before application of the pressure pulse and the current measured at the peak and the maximum of the fast and the slow components, respectively. The gray symbols indicate individual values. ^***^, = p < 0.001, paired-sample *t*-test; N = 2 different donors’ eyes, *n* = 15 and n = 9 cells for the fast and the slow components, respectively. **(F)** Bar graphs summarizing kinetics of fast and slow components. Results obtained from cells isolated from two different donors eyes were pooled. Shown is the mean ± SEM of time measured between application of the pressure pulse and when current reaches the maximum value. Grey symbols represent individual values. n = 17 cells and n = 6 cells for fast and slow response, respectively.

### Shear flow

Shear stress-induced changes in intracellular calcium were tracked in a microfluidic chamber designed for laminar flow, precise control of shear and full access to microscope objectives (Warner Instruments).

### F-actin and focal adhesion staining

TM cells were plated on type I collagen coated glass coverslips for 48 hour before undergoing treatment with Yoda1, GsMTx4 or vehicle for 1 hour at 37°C in cell culture CO_2_/O_2_ incubator. Slides contained fixed cells were probed with Phalloidin 488 (1:1000; Invitrogen), and antibodies raised against vinculin (1:1000; Sigma). Secondary antibodies were anti-mouse IgG Alexa Fluor 647 (1:1000; Invitrogen).

### Outflow Facility Measurements

C57BL/6 mice (2 males and 6 females, 3-6 months old) were euthanized by isoflurane inhalation followed by decapitation. Outflow facility was measured using the iPerfusion system, which is specifically designed to measure the low flow rates in the outflow system of paired mouse eyes (35).

### Data analysis

Student’s paired *t*-test or two-sample *t*-test was applied to estimate statistical significance of results. P < 0.05 was considered statistically significant. Results are presented as the mean ± S.E.M.

## Results

### SAC activity in TM cells involves kinetically separable components

We investigated mechanotransduction in primary TM cells isolated from two healthy donors. The cells expressed standard TM markers, including *MYOC, TIM3, AQP1, MGP* and *ACTA2* and responded to steroid DEX with MYOC upregulation **(Supplementary Fig. 1)** (34). Stretch-activated currents were measured under voltage clamp in the whole-cell configuration, at the holding potential of - 40 mV that approximates the cells’ resting potential (17). The cell membrane was expanded by application of hydrostatic pressure (1.5 sec, 25 mm Hg) delivered by high-speed pressure clamp (HSPC) **(Fig. 1A - C)**, a method that has been widely used to study SAC activation (36-38). Application of pressure steps evoked ionic currents in the majority (55/57; 96%) of recorded cells. A transient current (referred to as the “fast component”), seen in 72% of cells (n = 41/57), showed an average amplitude of -76.1 ± 11.3 pA/pF (mean ± S.E.M.) The pressure response reached peak amplitude within 1.18 ± 0.13 sec **(Fig. 1D - F)**.

The second component of the current induced by HSPC was characterized by delayed onset (peak response latency of 113 ± 8 sec), and slow return towards the baseline conductance. The average maximal amplitude of the “slow component” was -3.7 ± 1.0 pA/pF (n = 9 cells). This component was observed in 45% cells **(Fig. 1D, E & F)** whereas ∼14 % cells exhibited both components. These data demonstrate that TM cells isolated from healthy donors consist of functional subpopulations that differ by the kinetics of nonselective cation SAC conductances.

### Piezo1 mediates fast and TRPV4 mediates slow SAC activation in human TM cells

We investigated whether the time-dependence of the pressure-induced response involves distinct SACs. To identify the ion channel mediating the fast component, pressure steps were applied in the presence of Ruthenium Red (RR; 10 µM), a nonselective polycationic inhibitor of calcium-permeable mechanochannels. RR inhibited the fast component by 77 ± 7 % (n = 7 cells; P < 0.01) whereas HC067047 (5 µM), a selective blocker of TRPV4 channels had no effect (n = 15 cells, P > 0.05). However, pretreatment with the *Grammostola spatulata* mechanotoxin GsMTx4 (5 µM), a relatively selective extracellular blocker of the Piezo family (39, 40), attenuated the fast component from -76.1 ± 11.3 pA/pF to -19.6 + 5.7 pA/pF (∼86 %; n = 15 cells; P < 0.01) **(Fig. 2A & B)**. Piezo involvement was tested more precisely in cells overexpressing a Piezo1 shRNA construct that demonstrated 50-60% knockdown relative to scrambled Sc-shRNA (**Fig. 2C & D)**. The pressure response in in Sc-shRNA-transfected controls (−51.4 ± 9.9 pA/pF; n = 4 cells) was attenuated from to -5.5 ± 4.9 pA/pF (n = 4 cells) following treatment with Piezo1 shRNA, a ∼85% decrease (P < 0.05) **(Fig. 2E)**.

**Figure 2.**
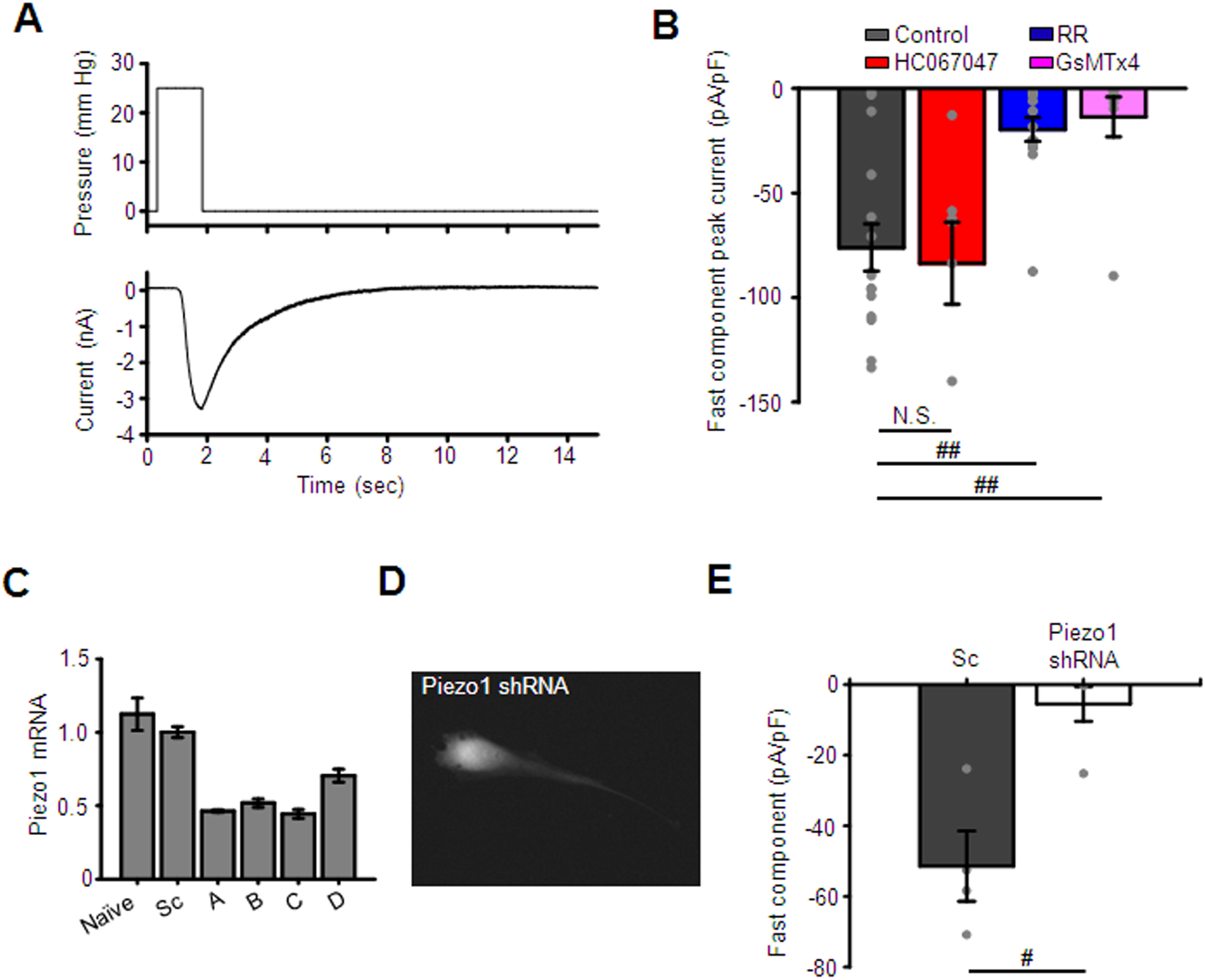
Piezo1 mediates the fast component of pressure-induced current. **(A)** A representative trace of the fast component. The waveform of the pressure pulse is shown above the trace. **(B)** The fast component is attenuated by non-selective Piezo1 antagonists ruthenium red (RR, 10 μM) and GsMTx4 (5 μM), but not a TRPV4 antagonist HC06047 (5 μM). Shown are the means ± SEM values of the fast response. The response was calculated by subtracting the baseline current from the peak current. ^##^, = p < 0.01, two-sample *t*-test. N = 2 eyes, n = 15, n = 7, n = 15, and n = 9 cells for untreated (control) cells and cells treated with HC067047, RR and GsMTx4, respectively. **(C)** Validation of Piezo1 shRNA constructs by Q-PCR analysis. Shown are the means ± SEM. **(D)** A representative fluorescent image of a TM cell overexpressing Piezo1 shRNA-mCherry construct. **(E)** Bar graphs summarizing effects of Piezo1 knockdown on the fast component of pressure response. The response was calculated by subtracting the baseline current from the peak current. Shown are the means ± SEM. ^#^, = p < 0.05, two-sample *t*-test. n = 4 cells and n = 4 cells for scrambled control (Sc-shRNA) and Piezo1-shRNA overexpressing cells, respectively.

To gain insight into channel kinetics, we measured single channel currents in excised patches of the TM membrane. Pressure steps (−80 mm Hg) induced events in 12/36 patches (**Fig. 3A**). In agreement with the reported conductance of human Piezo1 (41), the amplitude of the average unitary current was 2.53 ± 0.2 pA (n = 12; **Fig. 3B**), corresponding to an average conductance of 25.3 ± 2.0 pS. Mechanosensitive 25 pS events was not observed in patches from cells that overexpressed Piezo1 shRNA (0/27 patches) whereas Sc-shRNA-treated cells showed channel activity in 13/35 patches (**Fig. 3C**).

**Figure 3.**
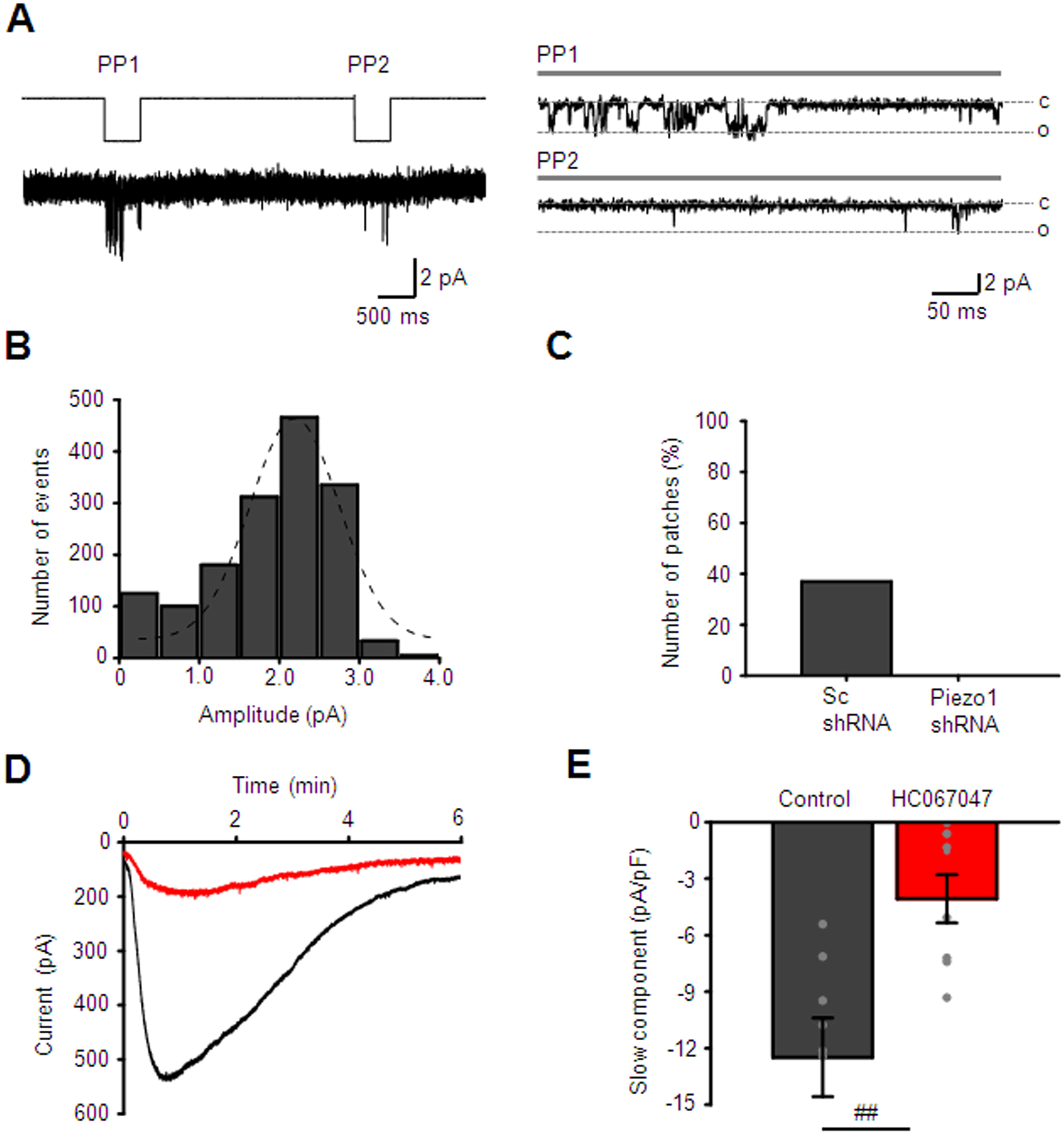
Activity of Piezo1-like channel in TM cells. **(A**) A duplet pressure pulse (−80 mm Hg, 500 ms) triggered the activity of a rapid activating and inactivating channel in excised inside-out plasma membrane patches. C: closed, O: open. **(B)** Amplitude histogram of channel opening events. **(C)** Occurrence of the Piezo1-like channel reduced in cells overexpressing Piezo1 shRNA compared to control cells overexpressing scramble shRNA. **(D)** Representative traces of the slow component illustrating the inhibitory effect of HC06047 (5 μM). The holding potential was -100 mV. **(E)** Summary for results illustrated in D. Shown is the means ± SEM of the fast response. The response was calculated by subtracting the baseline current from the peak current. ^#^, = p < 0.05, two-sample *t*-test. n = 8 and n = 8 for untreated (control) and HC067047-treated cells, respectively.

To get a more precise estimate of the time course of the fast component, we evaluated the latency of the indentation response (20, 42). Poking cells with a glass probe evoked robust inward currents with the activation and inactivation kinetics of 1.1 ± 0.2 and 1.8 ± 0.4 msec, respectively (**Supplementary Fig. 1**). The time course of the fast response indicates that the channel is directly activated by lipid distension (43-45). GsMtx4 suppressed 71.1 ± 9.2 % of the poke-induced current (**Supplementary Fig. 1**). These results suggest that the fast, inactivating response to pressure steps is largely mediated by Piezo1.

Given that TRPV4 is expressed in mouse and human TM, is activated by membrane strain and may modulate the outflow facility (15, 17), we wondered whether it contributes to the slow response component. At the holding potential of -100 mV, pressure steps evoked a robust, slowly-developing negative current in 8 out of 9 cells (−12.5 ± 2.1 pA/pF, n = 8 cells) that was sensitive to HC067047. The antagonist attenuated the peak current to -4.1 ± 2.1 pA/pF (n = 8 cells), with an average suppression of ∼73 ± 17% **(Fig. 3 D & E)**. Thus, SAC transduction in TM cells involves kinetically and functionally distinct Piezo1 and TRPV4 components.

### Molecular characterization of Piezo1 expression

To gain insight into the relative expression and localization of SACs, we used RT-qPCR and immunohistochemistry in cells expressing the standard markers αSMA, AQP1 and MYOC (**Fig. 4A**). The relative expression pattern was dominated by Piezo1 amplicons, which exceeded TRPV4 levels by ∼4-fold whereas Piezo2 expression was negligible (**Fig. 4A & B**). Western blots using a validated rabbit polyclonal antibody (46, 47) confirmed the expected Piezo1 protein band at ∼280 – 300 kDa (n = 2 donors; **Fig. 4C)**. The antibody labeled isolated cells, intact human TM and mouse TM tissue, with cultured cells showing cytosolic and membrane signals (**Fig. 4C**). Anterior segments isolated from normotensive donors showed prominent Piezo1-ir, which colocalized with TM markers α-SMA, collagen IV (**Fig. 4D**) and aquaporin 1 (not shown). Piezo1 localization to inflow and outflow pathways indicates that it is well placed to participate actively as a sensor of pressure-induced changes. Similar to human, Piezo1 was robustly expressed in TM in normotensive mice, with the signal observed in the TM, the ciliary body and cornea **(Supplementary Fig. 2A)**. In addition, Piezo1 expression was observed in nonpigmented cells of the ciliary body, ciliary muscle, stromal keratinocytes, and corneal epithelial cells. To validate antibody labeling, we examined Piezo1 expression in TM from Piezo1^P1-tdT^ mice, which express the fluorescent reporter (tdTomato) that was inserted in front of the Piezo1 stop codon in exon 51 (30). **Supplementary Fig. 2B** shows prominent signal within the TM, the ciliary body and cornea.

**Figure 4.**
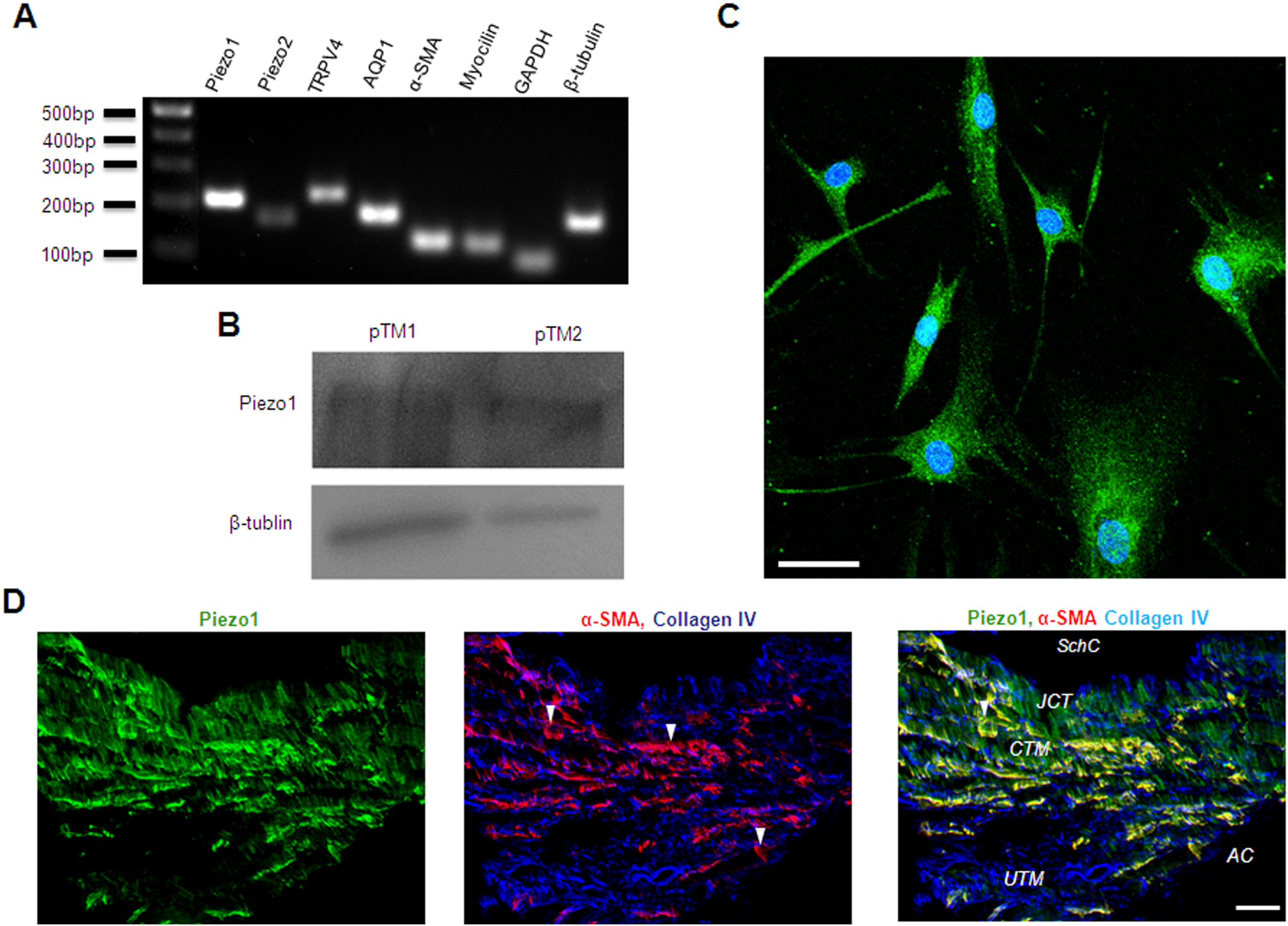
Molecular expression of Piezo1 channel in human TM. **(A)** Representative Q-PCR results demonstrating expression of mRNA encoding *PIEZO1, PIEZO2, TRPV4* and TM markers aquaporin 1 (AQP1), α smooth muscle actin (α-SMA) and Myocilin. **(B)** WB and **(C)** ICC results confirm the expression of Piezo1 protein in primary cultures of human TM cells. Scale bar is 50 μm. **(D)** Representative fluorescent IHC images illustrating the expression of Piezo1 (Green) in the anterior segment of the human eye. The tissue was co-stained for TM markers α-SMA (*red*) and Collagen IV (*blue*). SchC: the canal of Schlemm; TM: the trabecular meshwork. AC: Anterior chamber; JCT: Juxtacanalicular tissue; CTM: Corneoscleral trabecular meshwork; UTM: Uveal trabecular meshwork. Magnification bars: 20 µm.

### Piezo1 mediates nonselective cation influx and increase in [Ca^2+^]_i_

To verify that TM cells respond with Piezo1 activation in the absence of mechanical stressors, we assessed the response to the gating modifier Yoda1, which binds the C-terminal Agonist Transduction Motif (ATM) to stabilize the open pore (48, 49). Voltage-clamped cells responded to Yoda1 with an increase in the transmembrane current amplitude from -34.5 ± 7.8 pA at rest to -170.2 ± 39.6 pA **(Fig. 5A-C)**. The current-voltage (I-V) relationship of the whole-cell current was quasi-linear, showing a 32.6 ± 16.0 mV V_rev_ shift into the positive direction. The Yoda1-induced current inactivated in the continued presence of the activator **(Fig. 5A)**, with faster time course of inactivation at negative (−100 mV) than positive (+100 mV) holding potentials **(Fig. 5A)**. Analysis of Yoda1-induced changes in membrane potential showed a transient peak followed by sustained depolarization from the resting potential of -47.3 ± 4. 3 to -1.7 ± 2.2 mV. In all cells we observed transient hyperpolarizing component **(Supplementary Fig. 3A)**.

**Figure 5.**
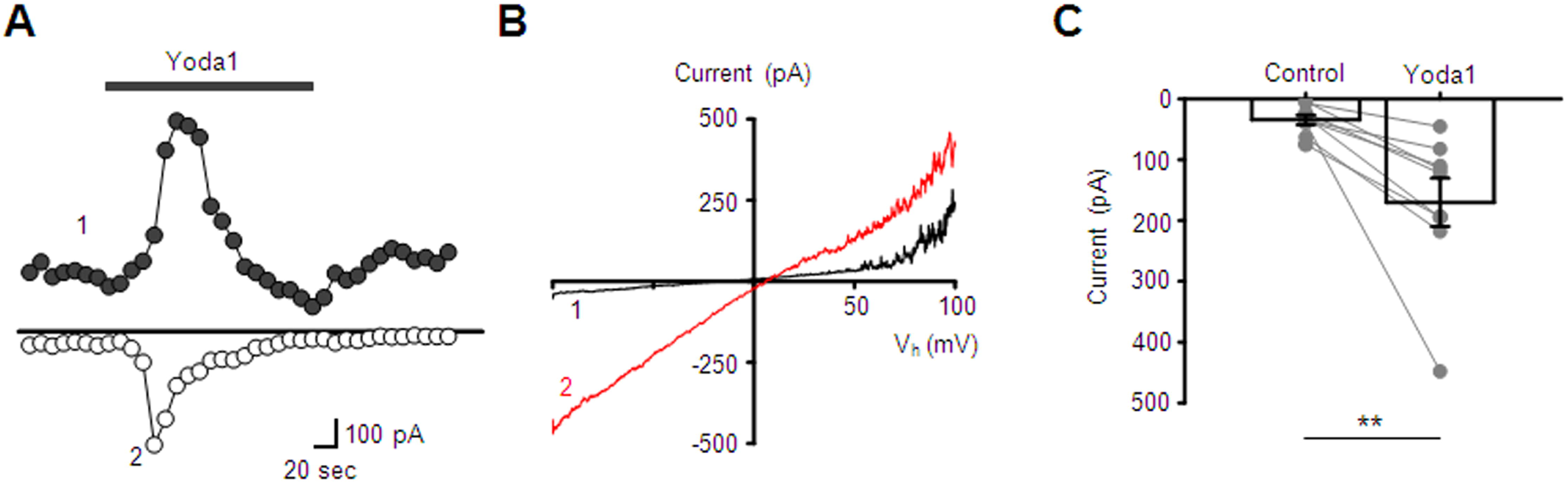
Agonist-induced activation of Piezo1 in TM cells. **(A)** Representative time course of the whole-cell current illustrating effect of Yoda1 (5 µM). Shown are the amplitude of current recorded at the holding potentials -100 mV *(open symbols)* and 100 mV *(filled symbols)*. **(B)** I-V curves of baseline (1, *black*) and Yoda1-evoked (2, *red*) current recorded at the corresponding time points in D. **(C)** Bar graphs summarizing effect of Yoda1 on the whole-cell current recorded at the holding potential -100 mV. ^*^, = p < 0.05, paired-sample *t*-test; N = 2 different donors eyes, n = 9 cells.

We assessed the spatial and temporal properties of Yoda1-induced Ca^2+^ signals, which might reflect the Piezo1 potential for regulating downstream signaling. The activator induced robust elevations in [Ca^2+^]_i_ across the TM cell (**Fig. 6A & C, Supplementary Fig. 3B**), which were abrogated by ≈ 99 % during the removal of extracellular Ca^2+^ **(Fig. 6 B & C)**. Ruthenium Red reduced Yoda1-evoked [Ca^2+^]_i_ signals by ≈ 68 % **(Fig. 6 D & E)** whereas GsMTx4 evinced ∼73 % inhibition (n = 17/15 for untreated control and GsMTx4-treated cells, respectively; P < 0.01) (**Fig. 6F & G**). Piezo1 knockdown with shRNA inhibited Yoda1-induced [Ca^2+^]_i_ signals by ∼50% (n = 25 and 17 cells for Sc and Piezo1 shRNA-treated cells, respectively; P < 0.05) **(Fig. 6H & I)**.

**Figure 6.**
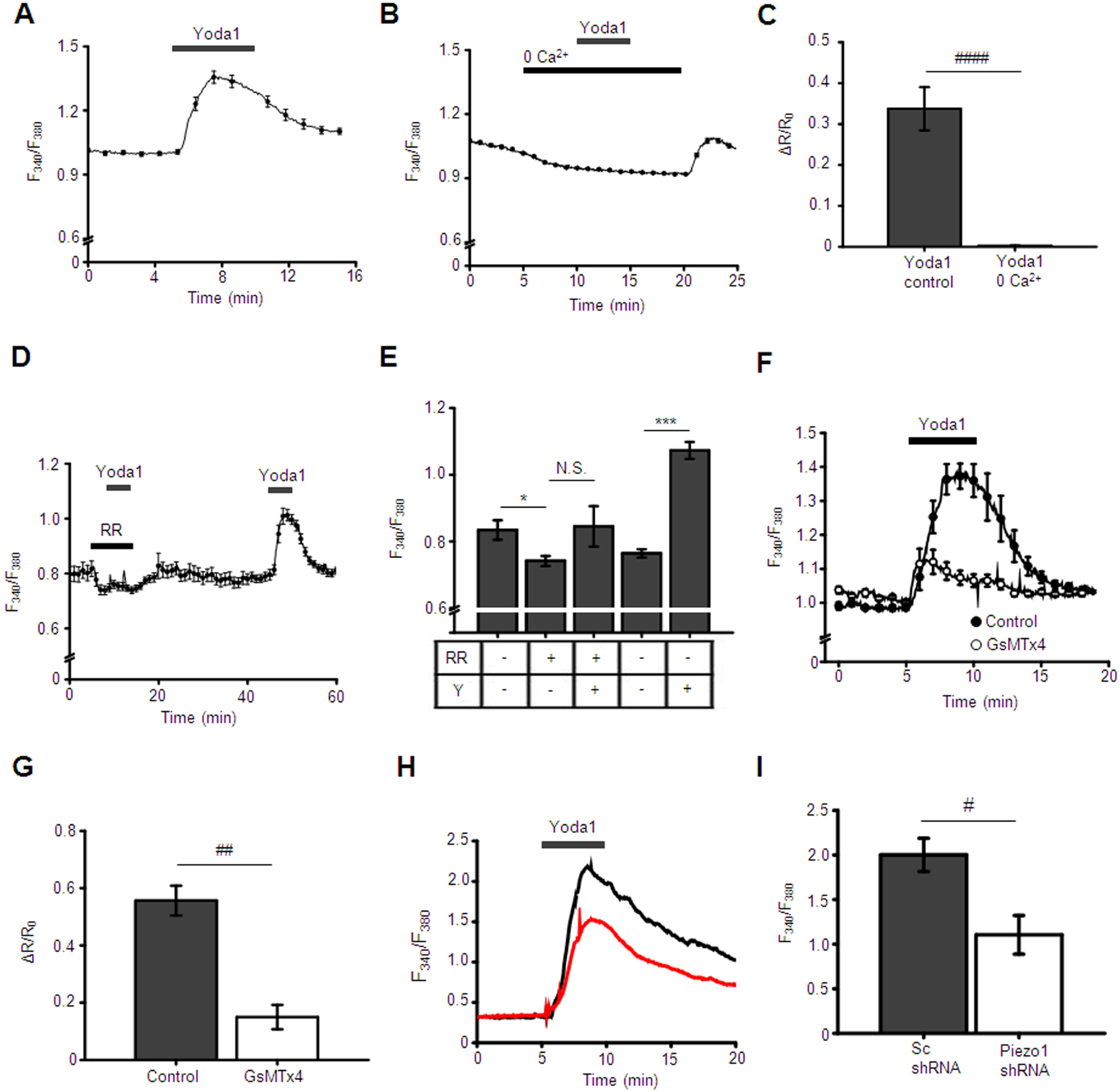
Activity of Piezo1 is functionally coupled to elevation of intracellular calcium ions in TM cells. **(A and B)** Yoda1 triggers a robust increase in [Ca^2+^]_i_ that was abolished in Ca^2+^-free extracellular solution. N = 2 eyes, n = 39 cells for control and N = 2, n = 61 cells for “0 Ca^2+^” conditions, respectively. **(C)** Bar graphs summarizing results illustrating in A and B. Shown the mean ± SEM. ^####^, = P < 0.0001; n > 50 cells, paired-sample *t*-test. **(D)** Ruthenium red (RR; 10 µM) abolished effect of Yoda1 (10 µM) on [Ca^2+^]_i._ **(E)** Bar graphs summarizing results shown in D. Shown the mean ± SEM. ^N.S.^, = P > 0.05, ^*^, = P < 0.05, ^**^, = P < 0.01; n = 14 cells, paired-sample *t*-test. RR: ruthenium red, Y: Yoda1. **(F)** Elevation of [Ca^2+^]_i_ by Yoda1 (10 μM) is attenuated in the presence of GsMTx4 (5 μM). Shown are averaged traces (the mean ± SEM) for untreated control cells (n = 17 cells; filled symbols) and GsMTx4-treated cells (n = 15 cells). **(G)** Bar graphs summarizing inhibitory effect of GsMTx4 on Yoda1-induced elevation of [Ca^2+^]_i_. **(H)** Representative F_340_/F_380_ nm traces illustrating effects of Piezo1 shRNA on Yoda1-mediated elevation of [Ca^2+^]_i_. The black trace represents control (Sc) and the red trace represents Piezo1 shRNA. **(I)** Bar graphs summarizing results shown in H. Shown the mean ± SEM. ^#^, = P < 0.05, ^**^, = P < 0.01; n > 50 cells, two-sample *t*-test.

Mechanical stimulation by cell poking with the glass probe induced a robust elevation of [Ca^2+^]_i_ by 296.9 ± 24.6 % relative to baseline levels. This effect was inhibited by GsMTx4 by 33 ± 8.8 **% (Supplementary Figure 4)**, indicating that nonselective cation currents and Ca^2+^ signals in TM cells are mediated by the mechanosensitive Piezo1 channel.

### Piezo1 is critically required for shear flow-induced TM [Ca^2+^]_i_ response

The bulk flow of aqueous humor driven by the pressure gradient imposes a drag force (shear) on the TM, particularly in the narrow passage ways of the JCT with predicted shears of 0.5 - 2 dynes/cm^2^ (50). To test whether TM cells are capable of responding to shear forces that mirror those encountered *in situ*, ratiometric Fura-2 calcium signals were measured in cells placed in a microfluidic chamber designed for laminar flow. The flow rate of 130 µL/min was applied through our shear chamber to produce a physiological shear stress of 0.5 dyn/cm^2^ (6). Exposure to shear elevated [Ca^2+^]_TM_ from baseline to the peak level of 0.821 ± 0.061 (n = 58), following which [Ca^2+^]_i_ recovered to a steady plateau. GsMTx4 reduced response amplitude by 86 ± 4.75 % (P < 0.004) (**Fig. 7B & C**), Ruthenium Red suppressed shear-evoked [Ca^2+^]_i_ signals by ∼45% (P < 0.01) whereas the TRPV4 antagonist HC067047 exerted only a slight inhibitory effect (17 ± 6.1%; P = 0.05) (**Fig. 7B & C**). These results identify Piezo1 as the principal transducer of shear flow in the TM.

**Figure 7.**
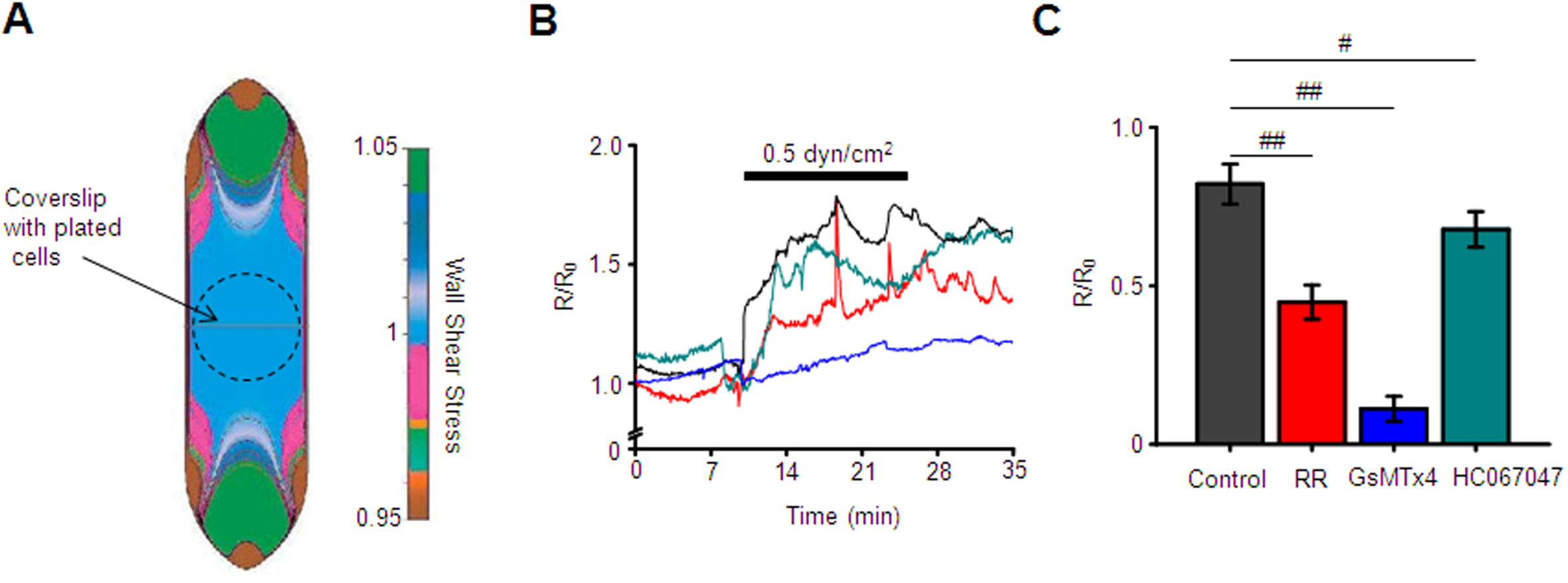
Piezo1 couples fluid shear stress to elevation of [Ca^2+^]_i_ in TM cells. **(A)** Distribution of wall shear stress in the flow chamber. The image was adapted from Warner Instruments. **(B)** Representative traces of normalized F_340/380_ ratio illustrating effect of laminar flow stress on [Ca^2+^]_i_. **(C)** Bar graphs summarizing results shown in B. ^#^ = p < 0.05, ^##^ = p < 0.01; n = 58 cells, n = 43 cells, n = 53 cells, and n = 47 cells for untreated (control), ruthenium red treated, GsMTx4 treated and HC067047-treated cells, respectively. Two-sample t-test. Shown are the mean ± SEM.

### Activation of Piezo1 upregulates F-actin and focal adhesions in TM cells

Actin cytoskeleton and focal adhesions are major determinants of TM stiffness and rigidity (51, 52), which in turn determine the outflow resistance (53) (19). Since activity of Piezo1 was implemented to remodeling of F-actin cytoskeleton and focal adhesions in various types of cells (54), we tested whether Piezo1 regulates remodeling of the cytoskeleton and focal adhesions in TM cells by assessing effects of Yoda1 on the immunoreactivity of F-actin and a focal adhesion protein vinculin. Treatment of TM cells with Yoda1 (2 μM and 10 μM) significantly upregulated F-actin (by 33.4 ± 4.1 % and 18.5 ± 4.8% by 2 μM and 10 μM of Yoda1, respectively) and increased the number of focal adhesions (41.2 ± 5.8 % and 62.8 ± 7.0 % by 2 μM and 10 μM of Yoda1, respectively) without significantly altering the cell area **(Fig. 8)**. We observed similar effect of Yoda1 on upregulation of phalloidin and focal adhesions using TM cells isolated from the eye of another healthy donor eye. These results indicate that activation of Piezo1 cause reorganization of cytoskeleton and focal adhesions in TM cells.

**Figure 8.**
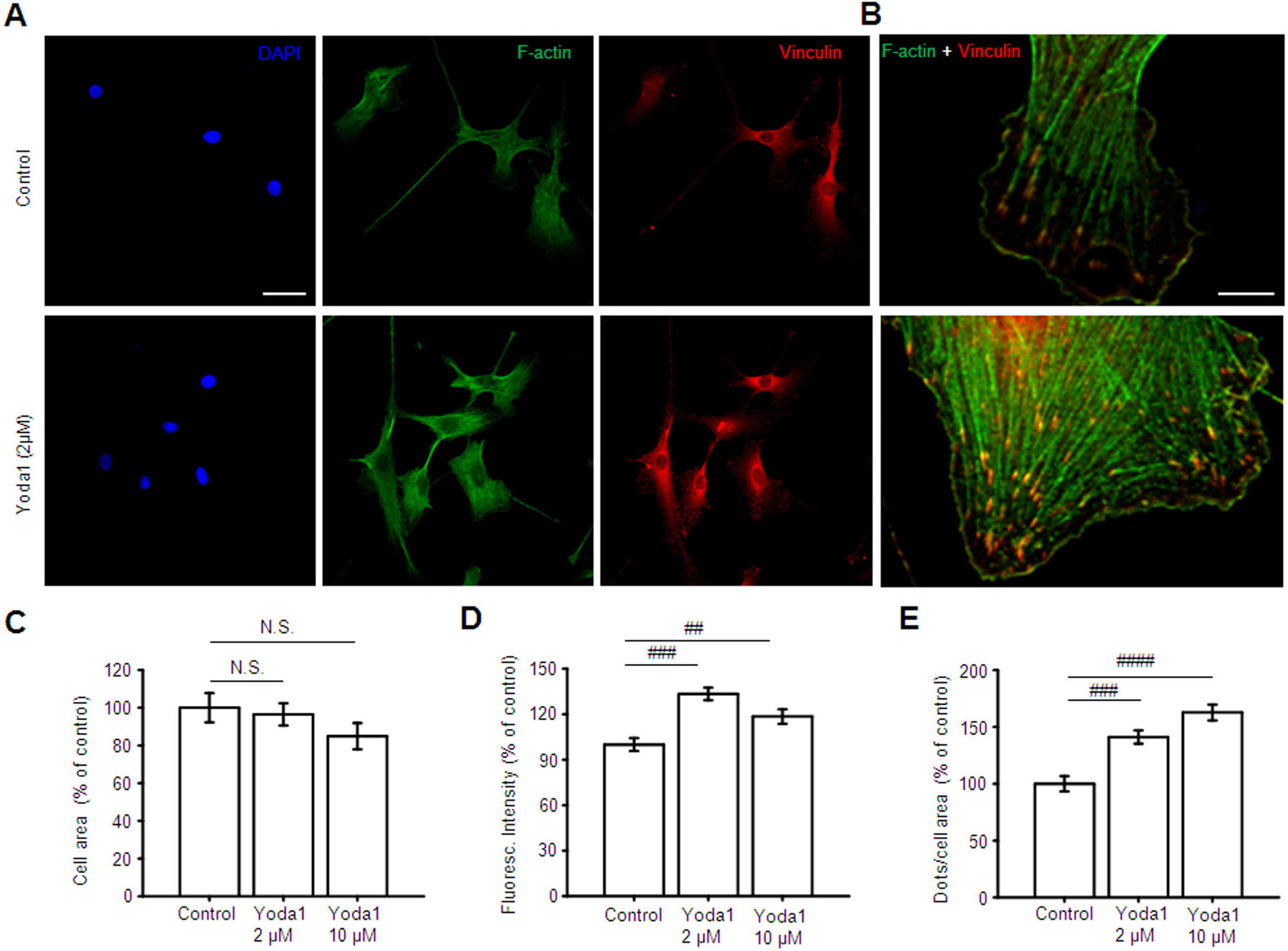
Piezo1 mediates reorganization of F-actin cytoskeleton and focal adhesions. **(A)** Representative examples of trabecular meshwork cells immunolabeled for F-actin (Phalloidin) and Vinculin with untreated (control) and Yoda1-treated TM cells. Yoda1 was applied at a concentration of 2 μM for 1h. Scale bar is 50 μm. **(B)** Magnified images of cells illustrating effects of Yoda1 on F-actin and vinculin. Scale bar is 10 μm. **(C – D)** Bar graphs summarizing effects of Yoda1 on the cells area, F-actin immunoreactivity and a number of vocal adhesions. ^N.S.^, = p > 0.05; ^#^, = p < 0.05; ^##^, = p < 0.01; ^###^, = p < 0.001; n = 41 cells, n = 65 cells, n = 59 cells, cells for control, 2 μM Yoda1 and 10 μM Yoda1-treated cells, respectively. Two-sample *t*-test. Shown are the mean ± SEM.

### Effect of PIEZO1 blockade on outflow facility

We sought to determine whether PIEZO1 activity is a component of pressure-induced regulation of TM-mediated hydraulic conductivity (“outflow facility”). Flow rate in response to sequential pressure steps was measured pairwise in enucleated mouse eyes, in the presence or absence of GsMTx4 (6 μM). The antagonist significantly reduced outflow facility (3.1 ± 0.3 nl/min/mmHg) as compared to control (4.6 ± 0.5 nl/min/mmHg, p < 0.01) **(Fig. 9)**, a 33% reduction. These data suggest that Piezo1 activity significantly augments fluid drainage via the pressure-regulated conventional outflow pathway.

**Figure 9.**
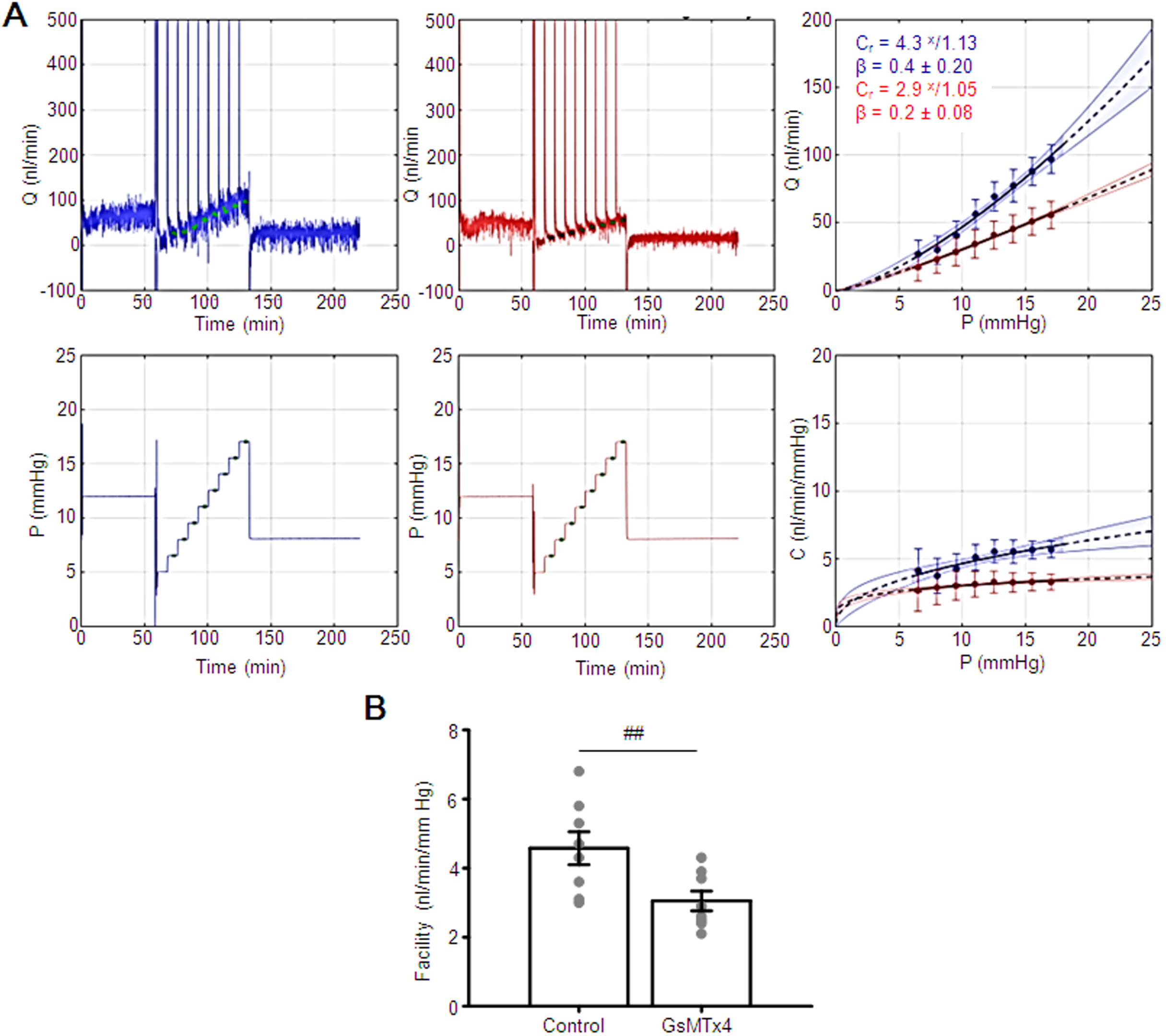
The activity of Piezo1 regulates outflow facility of trabecular outflow pathway. Shown are representative **(A)** traces that depict flow (Q) and pressure (P) measured in perfused, enucleated and paired mouse eyes. Traces show GsMTx4-treated *(red)* and untreated *(blue)* eyes. **(B)** GsMTx4 treatment reduced conventional, pressure-dependent outflow compared with no drug treatment (**p < 0.01, n = 8 eyes, ratio paired *t*-test).

## Discussion

In this study, we demonstrate that Piezo1 is required for the TM transduction of shear flow and membrane stretch, and link its activation to flow-induced cytoskeletal/focal adhesion remodeling. Piezo1 sensitivity to weak hydrodynamic loading, rapid activation and role in cell-ECM signaling support a model whereby the channel stabilizes and fine-tunes the outflow resistance in response to acute IOP displacements. Another, delayed SAC component was mediated by TRPV4 channels, which may collaborate with Piezo1 channels to impart mechanosensitivity to the ocular outflow system. The expression, sequence homology (24) and similar functional properties of Piezo1 in mouse and human TM suggest that its role in outflow regulation may be conserved across mammals.

It has long been obvious that the visual system is protected from pressure-induced neuropathy by sophisticated mechanotransduction mechanisms in the TM (4, 6). Putative mechanotransducers in TM cells include Piezo (55), TREK-1 (17, 56), TRPV4 channels (15), primary cilia (57), glycocalyx-actin interactions (58) and integrin-based adhesions (59). Similar to SMCs and vascular endothelial cells (14, 30), TM cells expressed high transcript levels of Piezo1 and TRPV4, but negligible levels of Piezo2. Western blots, double labeling with JCT-selective marker (αSMA), and analysis of tdT fluorescence from transgenic *Piezo1*^*tm1*.*1Apat*^ retinas confirmed Piezo1 expression in mouse and human TM. Accordingly, Yoda1, which stimulates Piezo1 but not Piezo2 (48, 49), evoked robust increases in [Ca^2+^]_TM_. Functional analyses that employed poking and pressure pulse steps revealed a fast mechanosensitive current with single channel conductance of ∼25 pS and strong inactivation. Suggesting that Piezo1 is required for TM mechanotransduction, the current was inhibited by Ruthenium Red, GsMTx4 and Piezo1-specific shRNAs. These findings are consistent with its functions in vascular pressure regulation and inflammatory mechanosensation (29, 46, 60). Its cognate Piezo2, previously linked to proprioception, somatosensation, cartilage remodeling and systemic baroreception (61-64) was localized to TM (55), however its low expression **(Fig. 4A)** argues against a major role in mechanotransduction.

In general, the levels of shear stress across the TM are believed to be negligible, except for the JCT region that may experience ∼0.5 - 2 dynes/cm^2^ and the Schlemm’s canal, where shear may reach up to 30 dynes/cm^2^ (6, 65). Given the negligible flow resistance, it has long been unclear whether shear stress constitutes a physiologically relevant stimulus for the TM (e.g., (6, 65). Our observation that shear flows 1-2 orders of magnitude lower compared to stresses typically used to stimulate endothelial and Piezo1-transfected HEK-293 cells (23, 30, 46) reliably produce calcium responses, suggests that aqueous hydrodynamics might be sufficient to regulate intracellular signaling in response to small IOP fluctuations that impose shear on juxtacanalicular cells (6, 66). The sensitivity of flow-induced signals to GsMTx4 and Piezo1 knockdown identifies Piezo1 as the principal transducer of these responses. The fraction (∼15%) of the flow-induced signal that was mediated by TRPV4 is in line with reports of TRPV4-dependent shear transduction in SMCs and endothelia (67-70). It is possible that the TRPV4 component might increase at shear stresses induced by larger pressure gradients (67-69).

It is not obvious how a rapidly inactivating channel (20, 71) mediates relatively sustained [Ca^2+^]_i_ signals in response to shear stress **(Fig. 7)**, however, low Piezo1 inactivation rates have been reported in cell-attached recordings from HEK-293 cells, chondrocytes, osteoblasts, endothelial and epithelial cells (46, 72-75). Importantly, sustained flow and pressure stimuli may remove Piezo1 inactivation without altering the pharmacology and unitary conductance of the Piezo1 mechanocurrent (23, 43, 74, 76). Potential inactivation-modulatory mechanisms include changes in membrane lipid composition, cytoskeletal modulation, amino acid alterations, sensitivity of N-terminal extracellular loop and the CED domains to recurrent force applications and/or presence of TMEM150c or ASIC1 proteins (73, 77-80).

Trabecular outflow is the principal IOP-regulating mechanism in rodent and primate eyes. The ∼30% reduction in pressure-dependent fluid outflow observed in GsMTx4-treated eyes **(Fig. 9)** suggests the possibility that Piezo1 activation resets the increase in pressure drop across the juxtacanalicular TM. The effect of the Piezo1 antagonist, which functions as an amphipathic channel blocker by lowering lipid strain on the channels (81) was comparable to glucocorticoids and TGFβ2, known drivers of ocular hypertension in glaucoma that reduce outflow facility by 23 - 46% (82-84) and 33%, respectively (85). The millisecond onset of Piezo1 activation **(Supplementary Fig. 1)** further suggests that Piezo1 can provide rapid, homeostatic adjustments in response to pressure steps.

Excessive actin polymerization, αSMA-dependent contractility and ECM upregulation represent key harbingers of increased outflow resistance in glaucoma (86, 87). Indicating that mechanotransduction contributes to use-dependent formation, alignment and plasticity of cell contacts, we found that exposure to Yoda1 suffices for the upregulation of stress fibers and vinculin-containing puncta. Similar results were observed in cells stimulated with TRPV4 agonists and pressure (15), pointing at calcium as the likely mediator of mechanically induced remodeling of actomyosin and cell-ECM contacts. Mechanosensitive Piezo1 and TRPV4 channels might therefore function as transducers of mechanical stresses within the local hydromechanical milieu and regulators of calcium- and use-dependent reorganization of TM structure and contractility. Taking into account the mechanosensitive TREK-1 channels that are likely to counterbalance pressure-dependent cation fluxes through Piezo1 and TRPV4 (16, 36), these data suggest that pressure homeostasis within the TM outflow involves at least three different mechanotransduction pathways.

The prominent effect of GsMTx4 on the outflow facility measured with iPerfusion implicates Piezo1 in the dynamic regulation of the primary outflow pathway. Under our experimental conditions, outflow principally reflected the conventional pathway. Future studies using conditional knockdown are needed to disambiguate the precise contribution of trabecular vs. endothelial stretch-activated channels within the canal of Schlemm (35), which together with the JCT TM layer contributes ∼75% of the outflow facility (88). In the intact animal, Piezo1 signaling in nonpigmented epithelial cells of the ciliary body and the ciliary muscle **(Supplementary Fig. 1B)** might provide additional functions in the regulation of fluid secretion and uveoscleral outflow.

Pressure steps applied to whole cell-clamped cells evoked an additional, delayed, current component. This current, mediated by TRPV4 channels, was not detected in excised patch recordings, presumably due to the absence of eicosanoid intermediates that are necessary for channel activation (15, 89). Its manifestation in the absence of the fast component and resistance to GsMTx4 argue against Piezo1-TRPV4 coupling seen in pancreatic acinar cells (70). Interestingly, the tissue resistance to aqueous outflow is augmented in response to Piezo1 blockade, TRPV4 agonists and GPCR-coupled store release, all of which modulate cytosolic [Ca^2+^]_TM_ (90). However, the following proof-of-principle examples suggest that functional divergence between Ca^2+^ pathways in nonexcitable cells is not uncommon: (i) Piezo1 promotes vasoconstriction in mesenteric endothelia whereas TRPV4 channels drives vasodilation (23, 91); (ii) Piezo1-mediated depolarization in SMCs activates voltage-operated calcium influx and vasoconstriction whereas TRPV4 -mediates vasodilation (92), and (iii) TRPV1 mediates contraction whereas TRPV4 mediates dilation of the ciliary muscle (93). We propose that Piezo1 and TRPV4 channels in TM cells are coupled to distinct microdomains and/or downstream Ca^2+^ effectors.

TM mechanotransduction plays an essential role in IOP regulation and glaucoma. Here, we show that Piezo1 is one of the mechanosensors that initiates the response to hydrostatic pressure and is required for the rapid response to pressure, stretch and shear flow, with possible functions in the regulation of aqueous outflow. Such “high-pass” activation (43) might sense IOP fluctuations impelled by ocular pulse, blinking, sneezing or yoga (94) to modulate pulsatile flow of the aqueous fluid (95) and protect the eye through time-dependent facilitation of trabecular outflow. Our data also reveal the collaboration between Piezo1 and TRPV4, a stretch-activated channel that mediates cytoskeletal and cell-ECM remodeling in the presence of chronic mechanical stress (15). It is possible that mechanosensory tuning of outflow resistance under different pressure regimens, segmental flows of aqueous humor, and tensile strains on trabecular lamellae, involves concurrent and balanced activations of multiple mechanosensitive channels that include Piezo1, TRPV4 and TREK-1 (15, 16, 96). Also worth noting are the many parallels with cardiovascular and pulmonary systems in which mechanochannel activation by fluid flow profoundly regulates hydrostatic pressure gradients associated with tissue development, function and pathology (26, 28, 29, 97, 98). The delineation of the intertwined mechanisms that mediate the TM sensitivity to mechanical stress may help identify novel potential targets that can be exploited for precise IOP control and stabilization in glaucoma.

## Acknowledgements

Supported by the National Institutes of Health (EY022076, EY027920, P30EY014800 to D.K. and NIH EY022359, NIH EY005722 to W.D.S.), Willard L. Eccles Foundation, The Neuroscience Initiative at the University of Utah and Unrestricted Grants from Research to Prevent Blindness to the Departments of Ophthalmology at the University of Utah and Duke University.

## Author contributions

O.Y., T.T.T.P., M.L.D., W.D.S. and D.K. designed research; O.Y., T.T.T.T.P., J.M.B., M.L.D, F.V.C., C.N.R., C.S. and M.L. performed research; O.Y., T.T.T.P., J.M.B., M.L.D, F.V.C., C.N.R., C.S., M.L., W.D.S. and D.K. analyzed data; O.Y. and D.K. wrote the paper.

