## Supplementary Information for "Mechanotransduction and dynamic outflow regulation in trabecular meshwork requires Piezo1 channels"

**Experimental Procedures**

*Animals.* C57BL/6J and *Piezo1^P1-tdT^* (*Piezo1^tm1.1Apat^*) mice were from JAX (#02914; Bar Harbor, ME). The initial transgenic 129/SvJ strain (1) was backcrossed to C57BL/6J for more than 8 generations. The animals were maintained in a pathogen-free facility with a 12-hour light/dark cycle and *ad libitum* access to food and water. Temperature was set at ~22-23˚C. No sex differences in the immunolabeling data were noted, so their data were pooled. Mice were 3 to 6 months in age. The studies were approved by the Institutional Animal Care and Use Committee (IACUC) of the University of Utah and followed recommendations of the NIH Guide for Care and Use of Laboratory Animals and the ARVO Statement for the Use of Animals in Ophthalmic and Vision Research.

*Cell culture and transfection*. TM cells were isolated from juxtacanalicular and corneoscleral regions of the human donors’ eyes, as described (2, 3), in accordance with consensus characterization recommendations (4). Passage 2 – 6 cells were seeded onto Collagen I-seeded coverslips and grown in Trabecular Meshwork Cell Medium (ScienCell, Catalog#6591) at 37^0^C and 5% CO_2_. Vehicle control scrambled shRNA (Sc-shRNA; Cat#: TR30021) and human Piezo1 shRNA (Cat#: TF313084) were purchased from OriGene. Cells were transiently transfected with shRNA constructs (5 μg per 25 cm^2^ tissue culture flask) using Lipofectamin 3000 reagent and utilized for experiments on 3^rd^-4^th^ day post-transfection. Transfected cells were recognized by expression of the fluorescent reporter (GFP and mCherry for Sc-shRNA and Piezo1 shRNA, respectively).

*Reagents.* Reagents used were largely purchased from Sigma-Aldrich. 2-[5-[[(2,6-Dichlorophenyl)methyl]thio]-1,3,4-thiadiazol-2-yl]-pyrazine (Yoda1) and *Grammostola spatulata* mechanotoxin 4 (GsMTx4) were from Sigma-Aldrich and Alomone Labs, respectively.

*Q-PCR.* Total RNA was extracted from TM cells using Applied Biosystems PicoPure RNA Isolation Kit (ThermoFisher Scientific) (5). 1 μg of total RNA was used for complementary DNA synthesis. RNA was reverse transcribed using qScript XLT cDNA Supermix (Quanta Biosciences). RT-PCR was performed using Apex qPCR 2x Master Mix (Genesee Scientific) with Applied Biosystems Veriti Thermal Cycler (ThermoFisher Scientific). PCR conditions were as follows: 95^o^C for 15min; 95^o^C for 20s, 60^o^C for 60s, 40 cycles. PCR products were run on a 2% agarose gel at 90V for 30min. The DNA bands were visualized by Ethidium Bromide staining along with 100-bp DNA ladder (IBI Scientific) using a FluorChemQ gel imaging system (Cell Biosciences).

*Western Blots.* Total protein samples were extracted from two primary trabecular meshwork lines from different patients in completed RIPA buffer supplemented with an enzyme inhibition cocktail (Biotechnology, Inc., Santa Cruz, CA, USA). Protein concentration was estimated with the Bradford assay. Total protein was heat-inactivated for 5 min at 95^0^C in Laemmli buffer (Bio-Rad Laboratories, Hercules, CA, USA) and allowed to cool for 15 minutes. Thirty micrograms of total protein per lane were loaded into 7% mini polyacrylamide gels. Electrophoresis was performed at 90 V for 15 min followed by 110V for 1.5 hr in the Mini-PROTEAN Tetra apparatus (Bio-Rad; running buffer 25mM Tris, 195 mM glycine, 0.1% SDS). Proteins were transferred to a polyvinylidene fluoride membrane (PVDF, 0.2uM, Bio-Rad Laboratories) in transfer buffer (25mM Tris, 195 mM glycine, 20% methanol) overnight at 4^0^C at 0.15 Amp. The membrane was blocked for 1 hour with 5% BSA in TBS-T, and incubated overnight at 4^0^C with an anti-Piezo1 antibody (1:200, Proteintech, Inc., Rosemont, IL, USA) or anti β-Tubulin (1:2000, Abcam, Plc., Cambridge, MA, USA) antibody. PVDF membranes were washed with TBS-T solution for 3 x 10 min and incubated in anti-rabbit-IgG HRP-linked antibody (1:2000, Cell Signaling Technology, Inc., Danvers, MA, USA) at room temperature for 2 hours. Protein bands were detected on X-ray film through enhanced chemiluminescence (ProSignal™ Pico; Genesee Scientific, Inc., San Diego, CA, USA) and developer (Merry X-Ray Corp., Menlor, OH, USA).

*Immunohistochemistry.* Anterior chambers were fixed in 4% para-formaldehyde for one hour, cryoprotected in 15 and 30% sucrose gradients, embedded in Tissue-Tek® O.C.T. (Sakura, 4583), and cryosectioned at 12 µm, as described (6) (7). The sections were probed with antibodies against Piezo1 (Proteintech, 15939), α-SMA (Sigma, A2457), and Collagen IV (EMD Millipore, AB769). Secondary antibodies included anti-rabbit IgG DyLight 488 (Invitrogen, 35552), anti-mouse IgG DyLight 594 (Invitrogen, 35511), and anti-goat IgG Alexa 647 (Invitrogen, A21469). Sections were coverslipped with DAPI-Fluoromount-G (EMS, 17984-24) and imaged with Fluoview-1000 confocal microscope (Olympus, Center Valley, PA).

*Calcium imaging.* TM cells were seeded on glass coverslip for 48 hours, loaded with Fura-2AM (Invitrogen) for 40-60 min, and washed with the bath solution containing (in mM): 140 NaCl, 4.7 KCl, 1.2 MgCl_2_, 5.6 glucose, 10 HEPES, 1.8 CaCl_2_ (pH 7.4, osmolarity 295-300 mOsm) for 5-30 min. Fluorescent imaging followed published protocols (8, 9). Excitation for 340 nm and 380 nm filters (Semrock, Rochester, NY) was delivered by a liquid light guide from a 150W Xenon arc lamp (DG4, Sutter Instruments). Fluorescence emission was high pass-filtered at 510 nm and captured with a cooled digital CCD camera (Photometrics) binned at 2x2. Data acquisition, F_340_/F_380_ ratio calculations and background subtraction were performed by NIS Elements 3.22 (Nikon) on Regions of Interest (ROI) encompassing the central cell area. In cell poking experiments, cells were loaded with Fluo-4 AM for 50 min at 37^0^C in CO_2_/O_2_ incubator. Poking of cells is described below. For data analysis, Fluo-4 fluorescence was normalized to mean baseline values obtained prior to poking.

*Electrophysiology.* Borosilicate patch pipettes (WPI) were pulled using a Flaming/Brown puller (Sutter Instruments), with resistance 6-8 MΩ when filled with the internal buffer solution containing (mM): 125 K-gluconate, 10 KCl, 1.5 MgCl_2_, 10 HEPES, 10 EGTA, pH 7.4. The chamber was superfused with saline containing (in mM): 140 NaCl, 2.5 KCl, 1.5 MgCl_2_, 1.5 CaCl_2_, 5.6 D-glucose, 10 HEPES (pH 7.4, adjusted with NaOH) (3, 10). The pipette solution in excised patch recordings was identical to the extracellular solution. Patch clamp data was acquired with a Multiclamp 700B amplifier, pClamp 10.6 software and Digidata 1440A interface (all from Molecular Devices). Data was sampled at 5 kH, digitized at 2 kH and analyzed with Clampfit 10.7 (Molecular Devices) and Origin 8 Pro (Origin lab). The holding potential in the whole-cell and inside-out excised patch recordings was set to -40 mV and 100 mV, respectively. In some voltage-clamp experiments, the holding potential was set to -100 mV to minimize contribution of TREK-1 channels (3).

Steps of positive (in whole-cell recording) and negative (in single-channel recording) pressure were delivered via High-Speed Pressure Clamp (ALA Scientific) as described (**Figure 1A)** (3, 10). In some experiments, membrane stretch caused by pressure application was visualized as increased fluorescence of the cell volume marker calcein **(Fig. 1B and C)** (11). The timing and intensity of pressure steps were controlled by Clampex software. To minimize membrane creep, membrane potential rundown and/or desensitization/inactivation (12, 13) a single stimulus was executed per cell in the whole-cell recording and two stimuli were applied to the excised patch. Single channel stimuli were -80 mm Hg and 500 msec duration, whole cell stimuli were -25 mm Hg and 1.5 sec duration. The single channel conductance was obtained through linear fit of the current-voltage plots. Current-voltage (I-V) curves of Yoda1-induced current were obtained from voltage Ramps ascending from the holding potential -100 mV to 100 mV (200 mV/sec).

Indentation responses were elicited with a glass pipette (tip diameter ~ 3 μm) position at a 30^o^ angle relative to the cover glass. The probe was positioned ~ 2 μm from the cell using a Sutter MPC-200 micromanipulator and indented with a rapid manual step to the depth of ~ 60 nm for 1 sec. The indentation step induced currents that varied by 8.2%. All experiments were performed at room temperature (20 – 22^0^C).

*Shear flow.* Shear stress-induced changes in intracellular calcium were tracked in a microfluidic chamber designed for laminar flow, precise control of shear and full access to microscope objectives (Warner Instruments). TM cells were plated on type I collagen coated glass coverslips 48 hours before being loaded into the shear flow chamber for experiments to prevent cell detachment due to shear flow. A programmable peristaltic pump (Harvard Apparatus) was used to control the total media flow rate, with shear Stress calculated using the following equation:

γ = V/x and Ʈ = γ*ƞ

with shear rate, γ, velocity of the moving layer, V, and distance between layers, x, shear stress, Ʈ, and viscosity, ƞ. γ experienced by cells in the chamber was estimated as 0.5 dyn/cm^2^. Fluorescence imaging was conducted with an inverted Nikon Ti microscope using a 40x objective.

*F-actin and focal adhesion staining.* TM cells were plated on type I collagen coated glass coverslips for 48 hour before undergoing treatment with Yoda1, GsMTx4 or vehicle for 1 hour at 37^o^C in cell culture CO_2_/O_2_ incubator. Cells were fixed in 4% PFA, washed in PBS, permeabilized with 0.1% Triton X-100, and exposed to the blocking solution (1% BSA, 0.3% Triton X-100/PBS) (2). Slides were probed with Phalloidin 488 (1:1000; Invitrogen), and antibodies raised against vinculin (1:1000; Sigma). Secondary antibodies were anti-mouse IgG Alexa Fluor 647 (1:1000; Invitrogen). DAPI-Fluoromount-G -coverslipped slides were imaged with a Fluoview-1000 confocal microscope (Olympus, Center Valley, PA). Images were processed with Photoshop CS6 (Adobe, San Jose, CA).

*Outflow Facility Measurements.* C57BL/6 mice (2 males and 6 females, 3-6 months old) were euthanized by isoflurane inhalation followed by decapitation. Eyes were immediately enucleated and used for experimental procedures. Each eye of pair was attached to a support platform in one of two identical perfusion chambers using a small amount of cyanoacrylate glue (Loctite, Westlake Ohio, USA). The perfusion chamber was filled with pre-warmed phosphate-buffered saline containing 5.5mM glucose and divalent cations (DBG), and temperature was maintained at 35°C. A glass microneedle was filled with DBG containing 6 μM GsMTx4 (Alomone Labs, Jerusalem, Israel) or DBG alone. The microneedle was connected to the perfusion system and was inserted into anterior chamber using a micromanipulator, while visualized under a stereomicroscope. Outflow facility was measured using the iPerfusion system, which is specifically designed to measure the low flow rates in the outflow system of paired mouse eyes (14). Initially, both eyes were perfused at 12 mmHg for 60 min to allow acclimation and delivery of the drug to cells of the outflow pathway, followed by 9 sequential pressure steps, with equal intervals between 5 and 17 mmHg. Data analysis was carried out as described previously (14). Briefly, a non-linear flow-pressure model was used to account for the pressure dependence of outflow facility in mice. A reference pressure of 8 mmHg was used to calculate the outflow facility. The order (left versus right and drug versus control) in which eye perfusions were performed was randomized.

*Data analysis.* Student’s paired *t*-test or two-sample *t*-test was applied to estimate statistical significance of results. P < 0.05 was considered statistically significant. Results are presented as the mean ± S.E.M.

**Supplementary Figure Legends**

**Supplementary Figure 1. (A)** A representative trace of the whole-cell current induced by cell poking in TM cells. Poking is indicated by an arrow. **(B)** The activation (τ_a_) and inactivation (τ_i_) time constants of poking-induced current. Time constants are represented as the mean ± S.E.M. **(C)** Current density. Shown are the mean ± S.E.M. ** = p < 0.01; two-sample *t*-test; n = 4 cells and n = 7 cells for control and GsMTx4, respectively. Gray symbols in the middle and the right plots represent individual values.

**Supplementary Figure 2. (A)** Expression of Piezo1 in the anterior segment of the mouse eye. Shown are representative IHC results of co-staining of the anterior segment with anti-Piezo1, anti-collagen IV and anti-α-SMA antibody. *aTM:* anterior TM, *pTM:* posterior TM, *SchC:* the Schlemm’s canal, *CB:* ciliary body. Scale bar is 50 µm. **(B)** The Piezo1^P1-tdTomato^ mouse from Jackson Labs (stock 029214) confirmed Piezo1 promoter activity in the trabecular meshwork and ciliary body. *aTM:* anterior TM, *pTM:* posterior TM, *SchC:* the Schlemm’s canal, *CB:* ciliary body. Scale bar is 50 µm.

**Supplementary Figure 3. (A**) Effects of Yoda1 on the plasma membrane potential of TM cells. The left panel: representative traces. Time of Yoda1 (10 µM) application is indicated by the bar. Arrowheads point at transient repolarizing component of the response to Yoda1. The right panel is a quantification of Yoda1-induced depolarization of the plasma membrane potential. Shown are the mean ± S.E.M. *** = p < 0.001; paired-sample *t*-test; n = 8 cells. Gray symbols represent individual values. **(B)** Yoda1 induced a robust elevation of [Ca^2+^]_i_ in TM cells. Shown are representative Fura-2 (F_340/380_ ratio) fluorescent images of cells taken before (control), during Yoda1 application and 10 min after washout of Yoda1.

**Supplementary Figure 4.** Cell poking induces elevation of [Ca^2+^]_i_ in TM cells. The left panel: averaged traces of Fluo-4 obtained from untreated (control) and cells treated with GsMTx4 (5 μM). The right panel: quantification of inhibitory effect GsMTx4 on poking-induced elevation of [Ca^2+^]_i_. Shown are the mean ± S.E.M. # = p < 0.05; two-samples *t*-test; n = 8 cells and n = 7 cells for control and Yoda1 plots, respectively. Gray symbols represent individual values.
