## Supplementary figures and images for "Mechanotransduction and dynamic outflow regulation in trabecular meshwork requires Piezo1 channels"

### Supplementary Figure 1

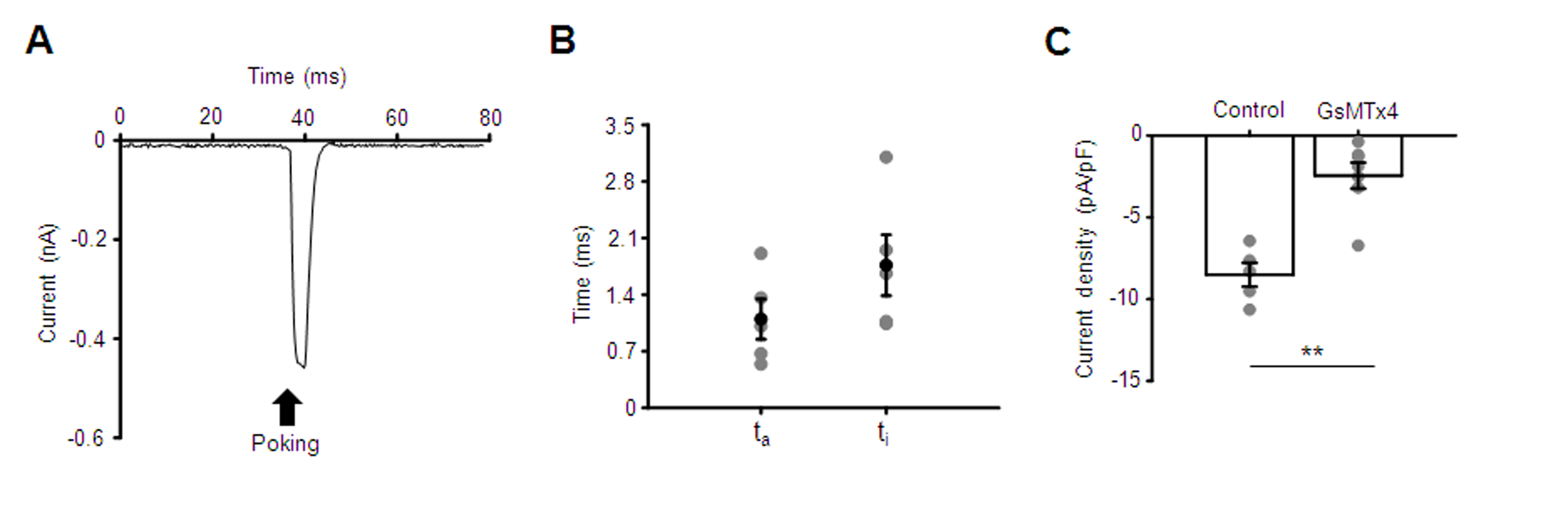

### Supplementary Figure 2

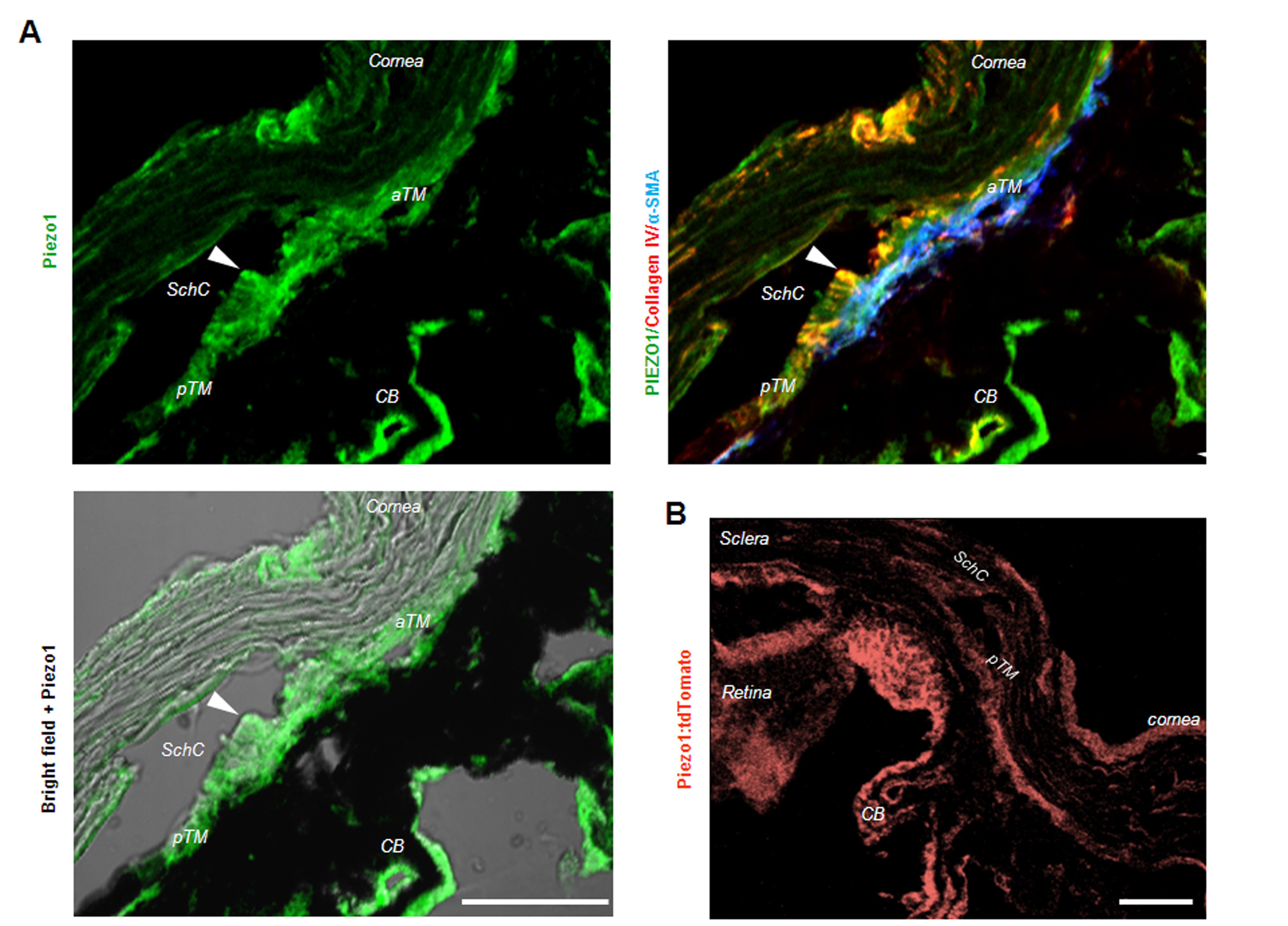

### Supplementary Figure 3

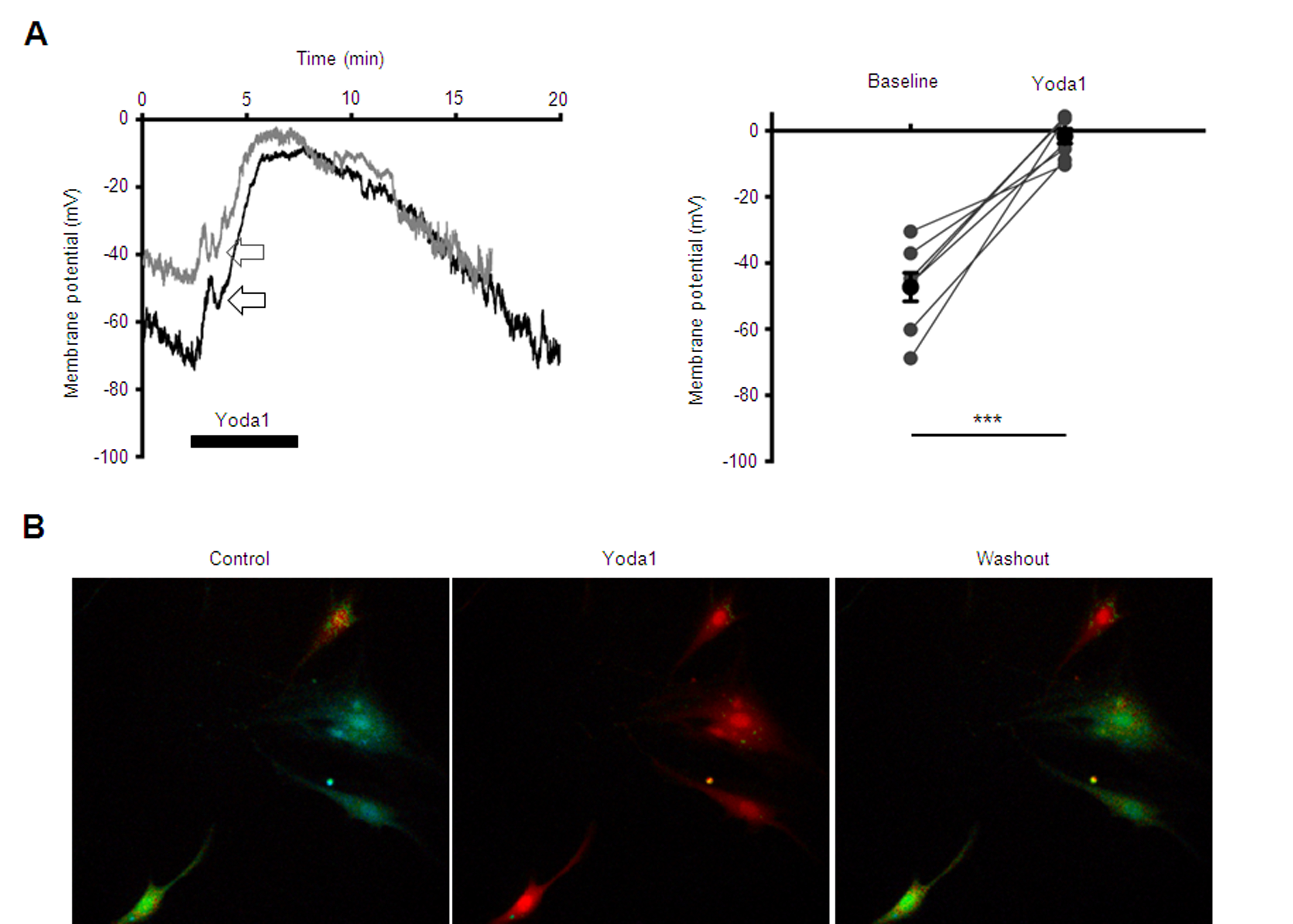

### Supplementary Figure 4

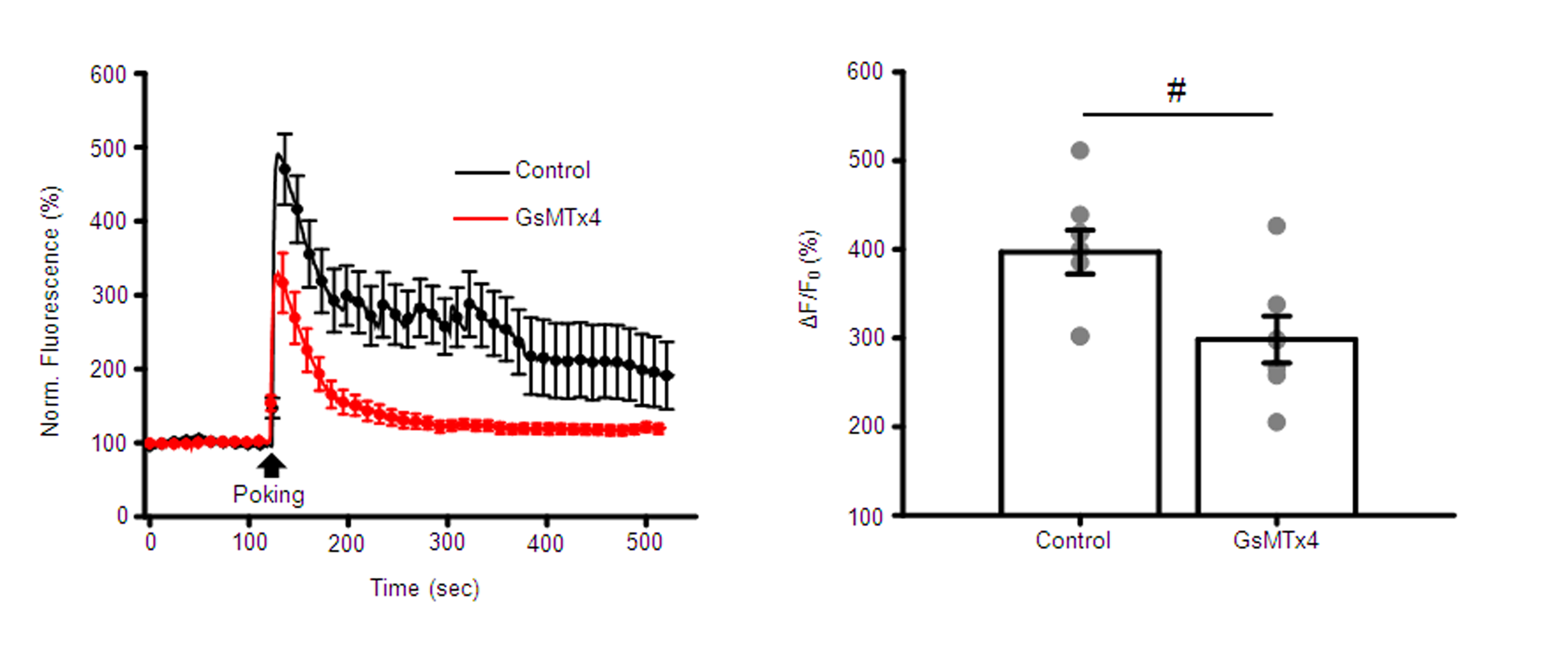
